# Hi-C deconvolution of a human gut microbiome yields high-quality draft genomes and reveals plasmid-genome interactions

**DOI:** 10.1101/198713

**Authors:** Maximilian O. Press, Andrew H. Wiser, Zev N. Kronenberg, Kyle W. Langford, Migun Shakya, Chien-Chi Lo, Kathryn A. Mueller, Shawn T. Sullivan, Patrick S. G. Chain, Ivan Liachko

## Abstract

The assembly of high-quality genomes from mixed microbial samples is a long-standing challenge in genomics and metagenomics. Here, we describe the application of ProxiMeta, a Hi-C-based metagenomic deconvolution method, to deconvolve a human fecal metagenome. This method uses the intra-cellular proximity signal captured by Hi-C as a direct indicator of which sequences originated in the same cell, enabling culture-free de novo deconvolution of mixed genomes without any reliance on a priori information. We show that ProxiMeta deconvolution provides results of markedly high accuracy and sensitivity, yielding 50 near-complete microbial genomes (many of which are novel) from a single fecal sample, out of 252 total genome clusters. ProxiMeta outperforms traditional contig binning at high-quality genome reconstruction. ProxiMeta shows particularly good performance in constructing high-quality genomes for diverse but poorly-characterized members of the human gut. We further use ProxiMeta to reconstruct genome plasmid content and sharing of plasmids among genomes—tasks that traditional binning methods usually fail to accomplish. Our findings suggest that Hi-C-based deconvolution can be useful to a variety of applications in genomics and metagenomics.

## Introduction

Microbiologists have pursued the characterization of the microbial flora of the human body since the 17th century (Lane, 2015). In recent years, this goal has been fulfilled in part by the development of genomic technologies that capture sequence information from highly complex consortia of microbial communities (Consortium, 2012; Qin et al., 2010; Turnbaugh et al., 2009; Schloss and Handelsman, 2005). The human gut microbiome, specifically, is of great interest both due to its role in basic human physiology and as a locus of microbial infectious disease. Sequencing of DNA directly extracted from mixed communities has allowed researchers to bypass the need for laborious isolation and culturing efforts for microbiological analysis. In addition to using such “metagenomes” to obtain insights about community-level dynamics and function, metagenomic data have also been used, with considerable success, for indirectly reconstructing genomes of individual species, both via *de novo* genome assembly (Zerbino and Birney, 2008; Peng et al., 2012; Nurk et al., 2017; Baker et al., 2010) and by statistical approaches for linking marker gene variation to genome content (Carr, Shen-Orr, and Borenstein, 2013).

However, while shotgun sequencing approaches have advanced, aided in part by the introduction of third-generation long-read sequencing technologies (Tsai et al., 2016) and new computational methods (Wood and Salzberg, 2014), our ability to accurately group DNA sequences or assembled sequence contigs into single-species genomes has remained limited. This grouping process, termed binning, often relies on a variety of indirect measures of consistency between contigs (Laczny et al., 2015; Wu et al., 2014; Imelfort et al., 2014; Dick et al., 2009). For example, contigs with similar read depth profiles are more likely to come from the same genome than those with vastly different read depths. Another source of evidence that two contigs come from the same genome is their tetra-nucleotide or k-mer frequency spectra; these signatures often delineate different bacterial species. Most methods use integrative binning methods, which result in more accurate genome grouping but require summarizing acrossmultiple samples (Kang et al., 2015; Wu, Simmons, and Singer, 2016). Moreover, binning is by nature an indirect clustering method, and sequence or read depth similarity does not necessarily imply that two contigs should be grouped into the same pseudogenome. Furthermore, mobile genetic elements (MGEs), such as plasmids, do not always share the copy number or sequence composition of their host genomes, and the same MGEs are often observed in different strains and species (Wu et al., 2014), rendering their binning especially challenging. These elements can only be assigned to their host genomes by physical separation, most commonly through culturing individual microbes, a task that is often difficult or impossible due to largely unknown microbial growth requirements.

An alternative approach for culture-free microbiome characterization relies on microfluidics methods, enabling the construction of genomes from sorted single cells (Rinke et al., 2013). This approach removes the need to deconvolute mixed groups of sequences and thus reduces the possibility of genome cross-contamination due to spurious associations between contigs. Single-cell sequencing, however, produces genomes that tend to show low completeness according to common criteria (Parks et al., 2015), requires complex instruments, and is prone to miss low-abundance organisms due to random sampling of single cells from a population.

A new and promising deconvolution technique for complex microbial communities employs Hi-C (Lieberman-Aiden et al., 2009; Burton et al., 2014; Beitel et al., 2014) (or a related method, 3C; Marbouty et al., 2017; Marbouty and Koszul, 2015). In Hi-C-based deconvolution, covalent linkages among DNA molecules in the same cell are induced by treating intact cells with formaldehyde, and these linkages are then ascertained with proximity-ligation and high-throughput sequencing. Using this technique, associations between contigs of the same genome are measured directly, largely obviating the need for complementary data. Furthermore, Hi-C proximity estimations represent an orthogonal data type and are easy to obtain at the scale of complex communities. This approach therefore combines the benefits of physical segregation of DNA molecules with the convenience and ease of traditional shotgun metagenomic sequencing. Additionally, by capturing inter-chromosomal junctions (such as plasmid-genome interactions) this method is capable of correctly grouping self-replicating mobile elements that violate assumptions of uniform copy number or composition with regard to their host genomes. Yet, considering the promise of Hi-C based deconvolution, to date, studies successfully applying this method to mixed populations are scarce and focus primarily on very simple or artificial communities (Burton et al., 2014; Beitel et al., 2014; Heil et al., 2017). Indeed, until recently, it was not clear, for example, whether Hi-C analysis could be scaled to complex microbial communities such as those inhabiting the human gut.

Here, we describe the application of ProxiMeta™, a Hi-C-enabled method developed by Phase Genomics Inc., towards deconvoluting the human gut microbiome. We demonstrate unparalleled performance in reconstructing individual genomes from this complex mixture compared to traditional shotgun metagenomic binning analysis. We also report 14 novel bacterial genomes discovered by applying ProxiMeta to a single fecal sample. Finally, we demonstrate the applicability of this approach for addressing biologically-and medically-relevant questions regarding the composition of the human gut community and the sharing of MGEs among microbes in the same community.

## Results

**ProxiMeta Hi-C deconvolution of a fecal sample.**

We applied ProxiMeta Hi-C (Figure 1) to a fecal sample from a healthy adult human as follows. First, we used shotgun metagenomic sequencing to obtain 151bp paired-end reads from this microbial community. We then assembled these reads using metaSPAdes (Nurk et al., 2017), generating 506,596 contigs containing a total of 677 Mb sequence with a contig N50 of 9,168 bp (see Methods and Table S1). In parallel, we generated paired-end Illumina Hi-C sequence reads from the same sample and mapped these to the shotgun assembly contigs using standard procedures (Methods). Combining the assembly and short-read mappings provides a network of Hi-C linkages between contigs, which serve as the input to a graph-based clustering algorithm. Resulting clusters smaller than 1000 bp were omitted, as is common in analyzing metagenomic assemblies (Wu et al., 2014). We treat the remaining clusters of contigs, which we refer to throughout as “genome clusters”, as putative genomes. The result is a set of sequences in FASTA format that we subject to downstream analysis with a suite of evaluative tools (Methods), most notably CheckM (Parks et al., 2015), which uses the presence of single-copy marker genes in the genome clusters to evaluate genome completeness and the overrepresentation of these genes to evaluate contamination due to improper clustering of multiple genomes. These are frequently-used measures for evaluating the quality of draft microbial genomes (Parks et al., 2015; Bowers et al., 2017), wherein high completeness and low contamination are desirable. In the rest of this report, we conduct a deeper analysis of these genome clusters, compared to the results of a parallel genome binning analysis, to validate their quality, plausibility, and usefulness for addressing fundamental questions concerning the partitioning of microbial genomes.

**Figure 1:**
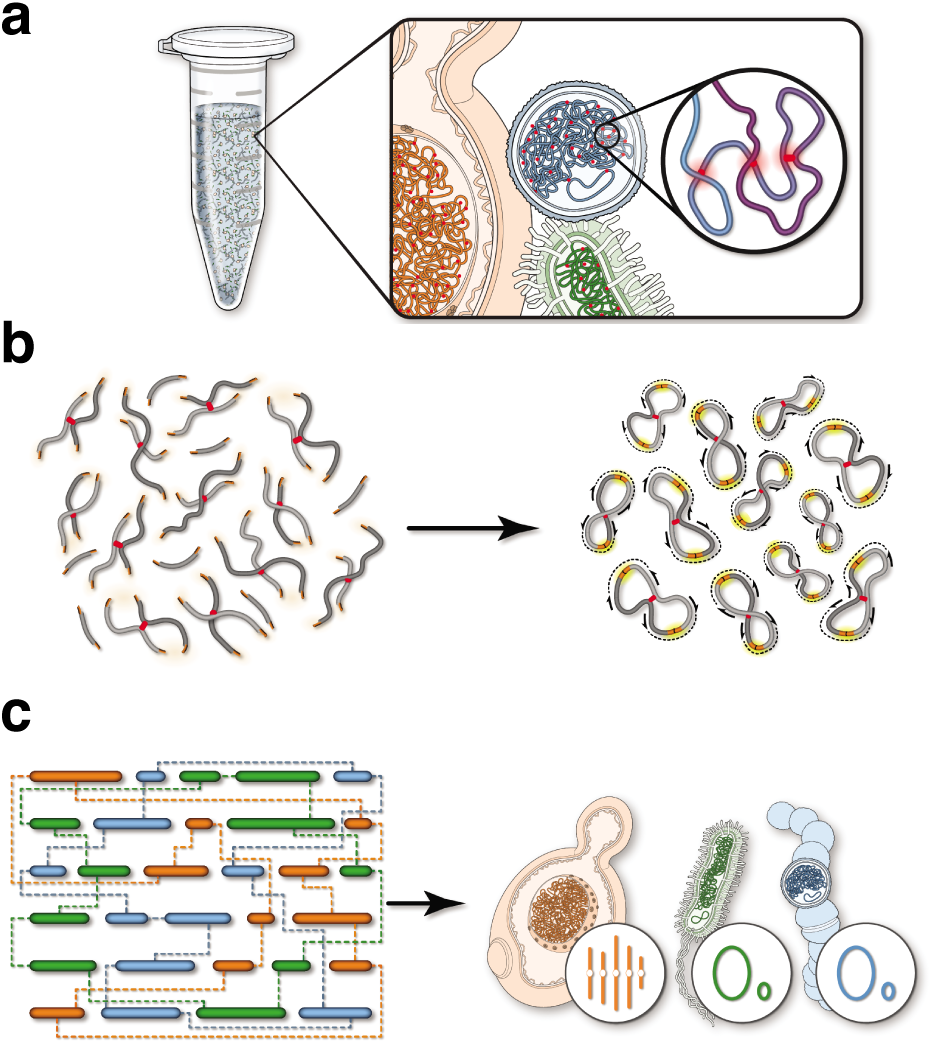
Schematic of the ProxiMeta method for metagenomic deconvolution. a) First, a sample consisting of mixed organisms is cross-linked with formaldehyde. Cross-links among DNA molecules are highlighted in red. b) DNA extraction yields a population of cross-linked DNAs with free ends from restriction cleavage. These free DNA ends are re-ligated and the resulting molecules read out with paired-end sequencing.c) Sequences ligated together and sequenced yield linkages between DNA contigs or scaffolds from an independently-generated sequence assembly. These linkages are used in clustering algorithms to deconvolute which DNAs derive from the same cell.

In total, ProxiMeta reconstructed 252 genome clusters from the fecal sample. From this set, 50 genome clusters were near-complete and 64 were substantially complete (<10% contamination for each; were greater than 90% complete more than 70% complete, respectively; Figure 2a, Table S2).

**Figure 2:**
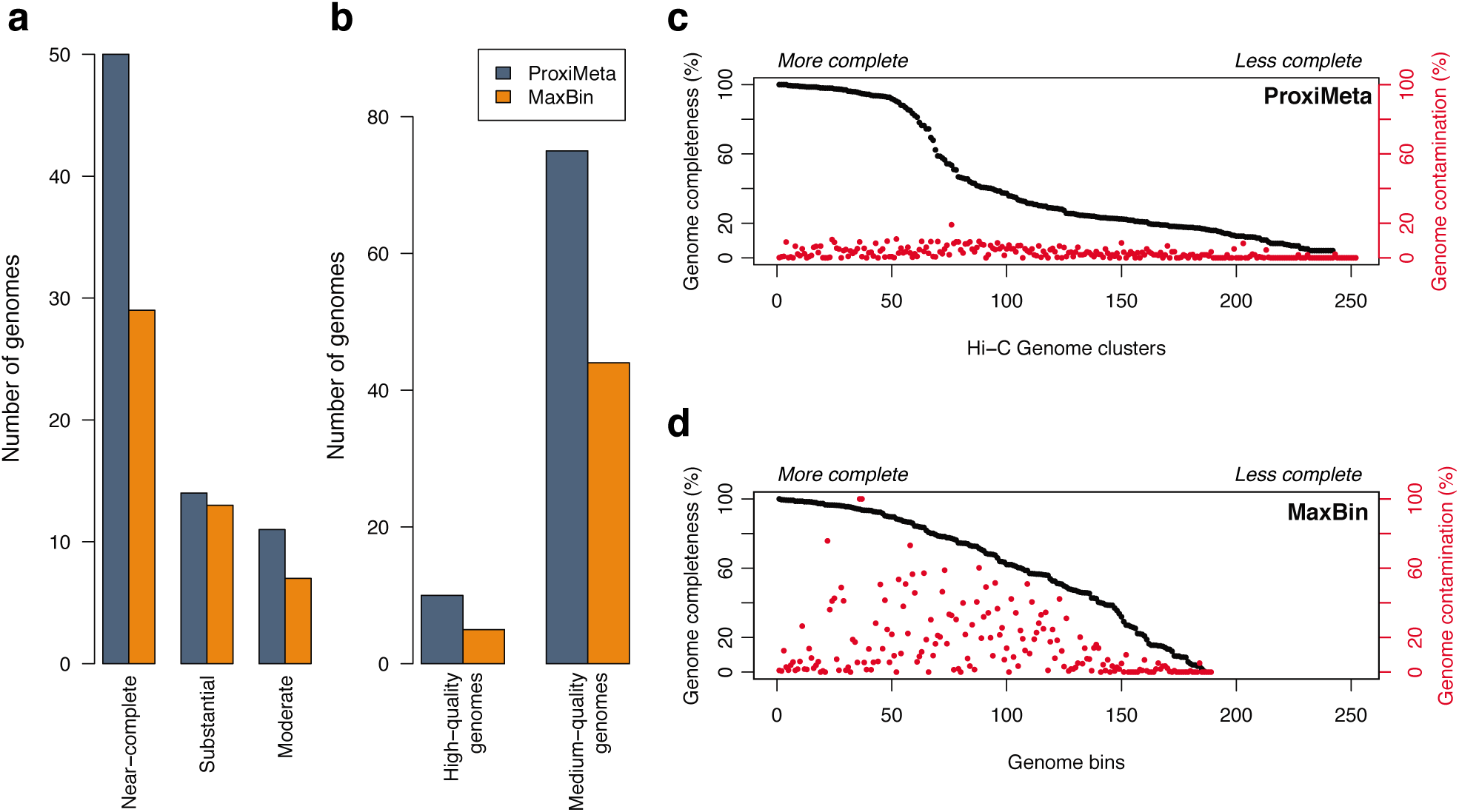
ProxiMeta Hi-C deconvolution recovers more genomes of higher quality from a single sample than genome binning procedures. a) Genomes adhering to CheckM criteria for completeness (near-complete: >90% complete, <10% contamination; substantially complete: >70% complete, <10% contamination; moderately complete: >50% complete, <10% contamination). b) Genomes adhering to MIMAG criteria for genome quality (high quality: >90% complete, <5% contamination, >=18 amino acids with tRNAs, 16S, 23S, and 5S rRNA genes present; moderate quality: >50% complete, <10% contamination). c) Completeness and contamination of ProxiMeta genome clusters, ordered by completeness. d) Completeness and contamination of MaxBin genome bins, ordered by completeness. Contamination values >100 were replaced with 100 for visualization purposes. (c) and (d) are scaled to have the same x-axis range.

These categories correspond to previously proposed thresholds for evaluating different levels of draft genome quality (Parks et al., 2015). Using a different set of criteria that also take into account rRNA and tRNA content, we found 10 “high quality” and 75 “medium-quality” draft genomes (Figure 2b). A total of 35,157 contigs >1000 bp in length were assigned to a genome cluster, which altogether accounted for 353 Mb of sequence. For further statistics on these clusters see Table S1 and Table S2.

## Comparison of ProxiMeta versus conventional metagenomic binning

To evaluate the performance of our ProxiMeta approach in comparison to more traditional binning methods, we used the same shotgun assembly contigs to infer genome bins using MaxBin, a commonly-used genome binning method based on coverage and tetranucleotide composition Wu et al., 2014; Wu, Simmons, and Singer, 2016. We compared the obtained genome bins to our Hi-C-assisted genome clusters. The MaxBin assembly yielded 29 complete or nearly-complete genomes (Figures 2a, 2b, Table S3). Additional criteria, such as the presence of universal RNA genes, indicated that genomes from the MaxBin assembly showed relatively high contamination relative to a Hi-C-based approach (Figure S2). There were 189 genome bins in total, composed of 89,622 contigs >1000 bp representing a total of 471 Mb of sequence. It is notable that the MaxBin output includes more total sequence than ProxiMeta, in spite of reconstructing fewer high quality genomes. ProxiMeta genome clusters collectively had an N50 twice as high as MaxBin genome bins (26.8 KB vs. 13.3 KB).

We next compared the results of the two procedures, ProxiMeta clustering and MaxBin binning, by examining whether they identify the same genomes. Given that we do not expect a perfect mapping of genomes, we used the MinHash k-mer hashing implemented in Mash (Ondov et al., 2016) to compute distances between each cluster/bin pair, and mapped MaxBin bins to ProxiMeta clusters if they were reciprocal best Mash hits with >500/1000 k-mer hashes as representing the same genome. By this criterion, among genomes >70% complete and <10% contamination, we found that the two methods reconstructed approximately the same genome in 32 cases, whereas MaxBin found 10 genomes missed by ProxiMeta, and ProxiMeta found 32 genomes missed by MaxBin. Thus, differences between the two methods are not primarily due to sampling different sets of genomes, but rather due to differing abilities to ascertain the same pool of genomes.

We compared the clustering/binning results to a null model, where contigs were shuffled randomly while preserving the size distributions of clusters and bins (Figure S3). We found that these shuffled groupings showed contamination roughly proportional to the total amount of sequence, unlike either method, and moreover relatively few random genome groupings showed high completeness under either method. Consequently, both ProxiMeta and MaxBin reconstruct genomes that are more plausible (more complete and less contaminated) than random contig groupings.

**Figure 3:**
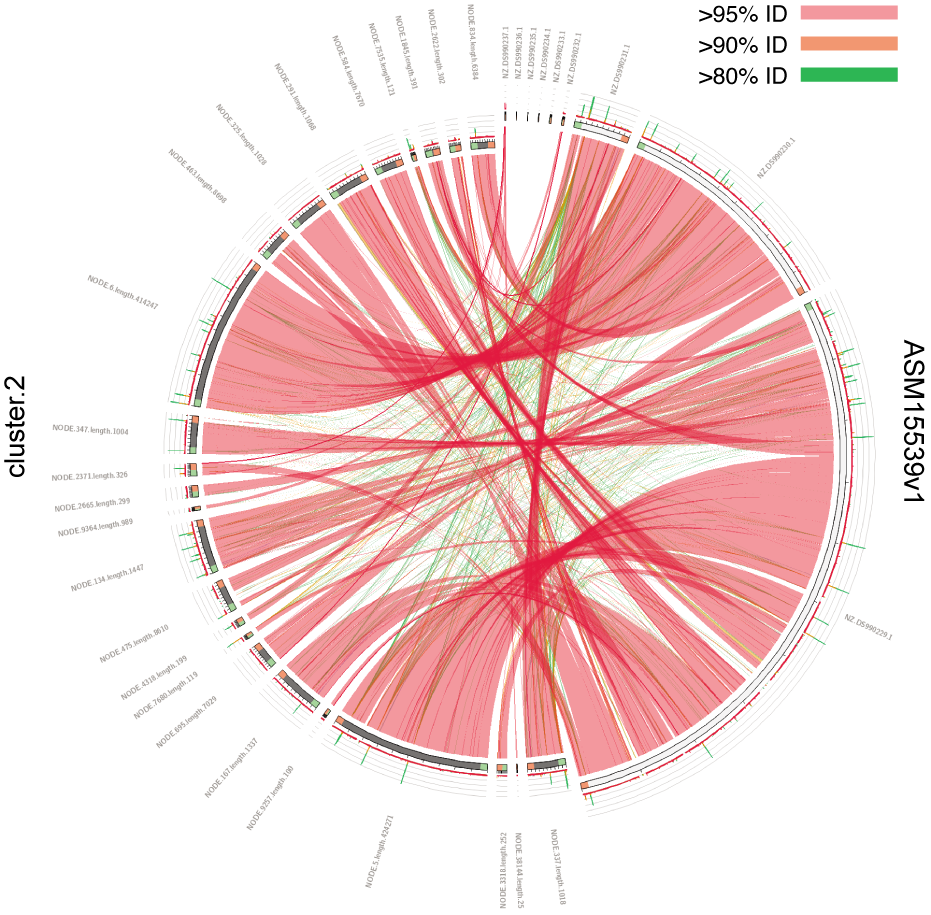
Representative genome from ProxiMeta deconvolution of a fecal sample align almost perfectly to known reference genomes. Each link drawn represents a BLAST alignment between the ProxiMeta genome cluster and the reference genome, with color indicating the percent identity of each pair of aligned sequence.

## Performance improvements of ProxiMeta are due to higher stringency of ProxiMeta genome clusters

As noted above, the differences between ProxiMeta and MaxBin were most marked for the highest-quality categories, in which ProxiMeta produces more genomes (Figure 2a, 2b). One explanation for the high quality of Hi-C genomes may be found by examining the contamination levels of the two sets of genomes. We found that ProxiMeta minimizes contamination compared to MaxBin (Figure 2c-d), suggesting that ProxiMeta correctly groups contigs based on evidence of physical proximity whereas MaxBin erroneously joins contigs from multiple genomes (presumably due to spuriously similar coverage and k-mer profiles). Common criteria for describing genomes require contamination <10%, which removes 43% of MaxBin genome bins from consideration, as opposed to 2% of ProxiMeta genomes.

It is also interesting to note that ProxiMeta produced many more low-contamination, low completeness genome clusters compared to MaxBin (Figures 2c, 2d). For example, the number of genome clusters with <50% completeness but also <10% contamination was 135 for ProxiMeta, but only 42 for MaxBin. This is further reflected in the overall smaller sizes of the ProxiMeta-derived genomes (Figure S4). This specificity likely reflects ProxiMeta’s use of DNA fragment proximity to avoid spurious contig groupings, yielding incomplete clusters of contigs but generally avoiding contamination by erroneously grouping contigs that in reality originated from different genomes. While these genome clusters are not complete characterizations of microbial genomes, they nonetheless contain valuable information about the community compared to ungrouped contigs.

We next assessed the relationship between quality and relative abundance of putative genomes from the two methods. Sequencing coverage of contigs is an important source of information for MaxBin (as well as for similar genome binning methods; Imelfort et al., 2014; Albertsen et al., 2013), but is not used by ProxiMeta, and thus the relative abundance of a genome in the community may be a contributing factor to performance differences. We observed that putative genomes from the two methods showed fairly similar relative abundances in range and overall distribution, though ProxiMeta genome clusters were somewhat less abundant (Figures S5a, S5b, p = 2.4E-6, Kolmogorov-Smirnov test). However, when we examined only fairly-complete genomes (>80% complete, <10% contamination) we found that fairly-complete genome abundances were similar between the two methods (Figures S5c, S5d, p = 0.20, Kolmogorov-Smirnov test). In both methods, the least abundant complete genomes have a measured relative abundance on the order of 0.05%. This analysis suggests that ProxiMeta’s superior performance is not strictly due to improved ascertainment of lower-abundance genomes, but due to more accurate ascertainment of moderate-to-high-abundance genomes. This was further confirmed by analyzing the relationship between abundance, completeness, and contamination across the two sets of putative genomes (Figure S6).

## Genome-genome alignment confirms that ProxiMeta faithfully reconstructs previously observed genomes

We investigated 5 genome clusters showing highest similarity to database genomes to confirm that genome clusters are plausible as draft genomes (Table 1, Figure 3, Figure S7, Table S4). We selected these 5 genome clusters based on >99% completeness, <2% contamination, and >600/1000 k-mer hashes matching their reference (Table S2). This approach of searching k-mer MinHash databases with ProxiMeta clusters tended to yield a single best hit (Table S2). These genomes generated contiguous and high-identity alignments to reference genomes, on average covering 93.3 +/-1.1% (Table 1). They also included relatively little additional sequence (93.3 +/-82.7 Kb) that did not align to the reference. Thus, for the “positive control” of known genomes, ProxiMeta generally reconstructs genome clusters consistent with references.

A similar analysis with 5 MaxBin genome bins revealed similar results (unsurprisingly, given that both represented the top genomes for their method), though these showed on average somewhat lower coverage of the reference (Table S5).

**Table 1:** Comparing reconstructed genomes to closely-related references.

| Genome | Assembly (cluster id) | Cluster length (Kbp) | Genome length (Kbp) | Genome fraction (%) | Percent identity (%) | Alignment Length (Kbp) |
| --- | --- | --- | --- | --- | --- | --- |
| ASM15539v1 | 2 | 2,137 | 2,203 | 92.5 | 93.0 | 2,039 |
| ASM98033v1 | 3 | 2,781 | 2,654 | 97.4 | 99.7 | 2,582 |
| MGS230 | 7 | 2,756 | 2,682 | 93.6 | 92.3 | 2,476 |
| MGS131 | 4 | 3,036 | 2,836 | 91.6 | 99.0 | 2,596 |
| MGS45 | 12 | 2,643 | 2,537 | 91.3 | 93.1 | 2,318 |

## Comparing taxonomic diversity ascertained using ProxiMeta vs. MaxBin

Interestingly, most of the genomes found by ProxiMeta but not by MaxBin were from the order *Clostridiales* (Figure 4a; 14 more genomes at level *Clostridiales*, 6 more genomes at level *Lachnospiraceae*). This suggests that ProxiMeta may be more successful at separating multiple related genomes than MaxBin or similar binning methods.

We visualized the taxonomic distribution of ProxiMeta genome clusters according to their degree of novelty, using single-copy marker genes to place them into a reference phylogeny. We found that both methods recovered genomes from substantially similar clades (Figure 4b, Figure S8), likely due to the restricted taxonomic range of the human gut microbiome. In agreement with the coarser last-common-ancestor analysis, we observed that ProxiMeta genome clusters without near matches in the RefSeq reference database were more frequently placed as Clostridial taxa, and were additionally quite widely distributed in this diverse but poorly-characterized taxon.

**Figure 4:**
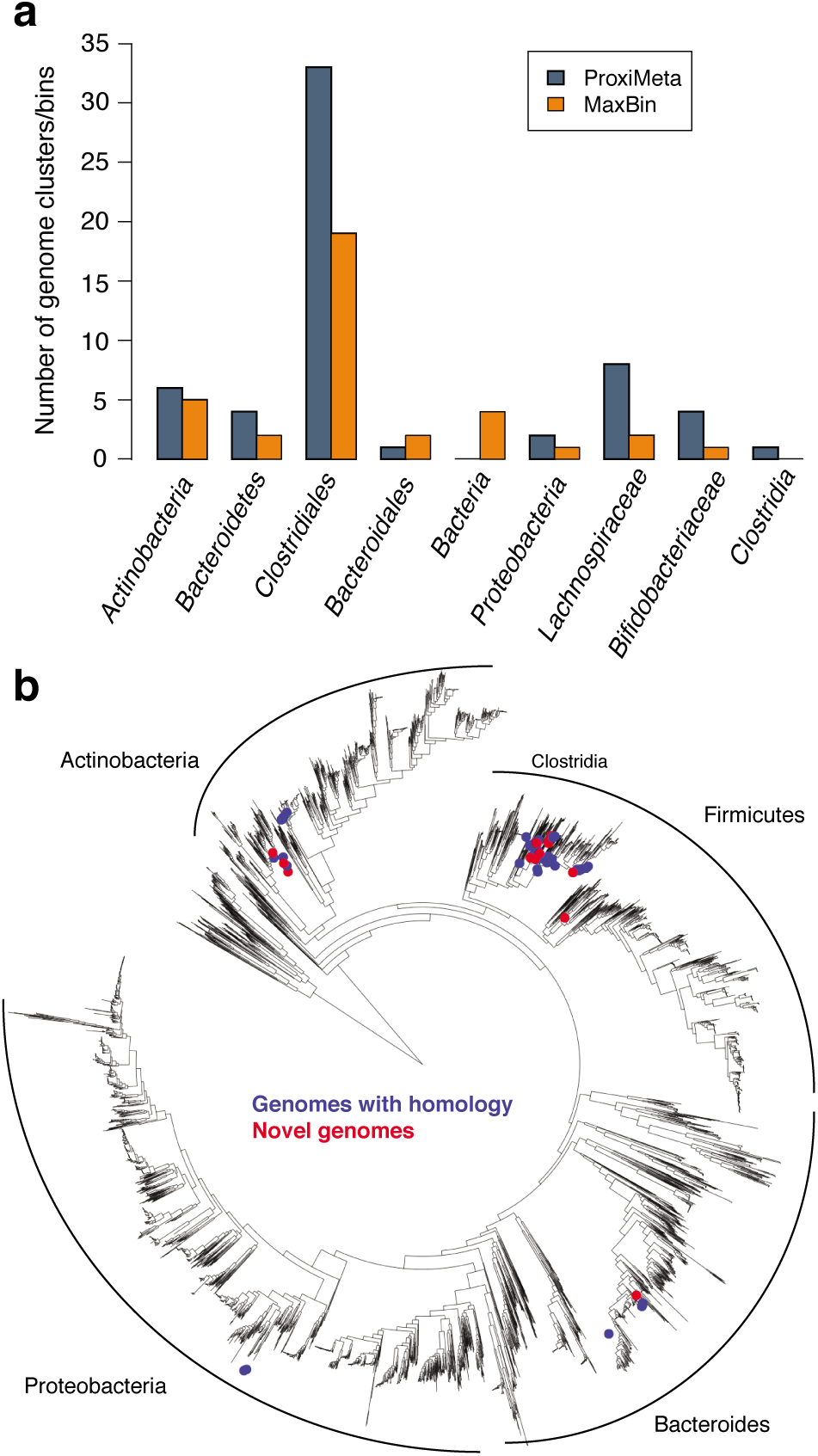
Phylogenetic placement of genomes derived from ProxiMeta. (a) All genomes included had completeness >80% and contamination <10%. The most specific possible taxonomic level was inferred using pplacer (Matsen, Kodner, and Armbrust, 2010) as run within CheckM on single-copy marker genes. (b) The tree is a prokaryotic phylogeny from Parks et al., 2015. Locations of genome clusters in the phylogeny are as determined by pplacer. Clades without any representation among genomes from this study were collapsed.

## ProxiMeta deconvolutes novel genomes

Although most of the genomes we recovered showed substantial homology to previously-described genomes, we also found 14 genomes >80% complete that appeared to be entirely novel (<20/1000 k-mer hashes to any single reference genome). Among these novel genomes, 10 belonged to class *Clostridia* (Figure 4b, Table S2), which is common in the gut and whose diversity is relatively poorly-characterized by culture-based methods (Manson, Rauch, and Gilmore, 2008; Eckburg et al., 2005). However, there were also prominent non-Clostridial novel genomes, for instance an actinobacterial genome of very high quality (cluster 1, Table S2).

## ProxiMeta accurately clusters plasmid contigs with genomes of origin

One of the most difficult challenges in metagenomic analysis is the ascertainment of plasmids and the identification of the organisms that carry them (Wu et al., 2014). ProxiMeta Hi-C’s intra-cellular proximity-ligation junctions capture inter-chromosomal DNA interactions and plasmid-genome interactions (Burton et al., 2014), and can therefore facilitate metagenomic-based plasmid analysis. To this end, we used a database of plasmid sequences derived from NCBI to identify putative plasmid sequences among all contigs in the original shotgun metagenome assembly, identifying in total 435 contigs with a >95% identity match of at least 500 bp to a plasmid sequence. Of these contigs, 185 showed at least one Hi-C read to be mapped. Among these 185 contigs, a majority (137) were assigned to a genome cluster (Figure 5a). Unclustered contigs were linked only to themselves, suggesting that intra-plasmid Hi-C interactions are stronger than plasmid-genome interactions.

**Figure 5:**
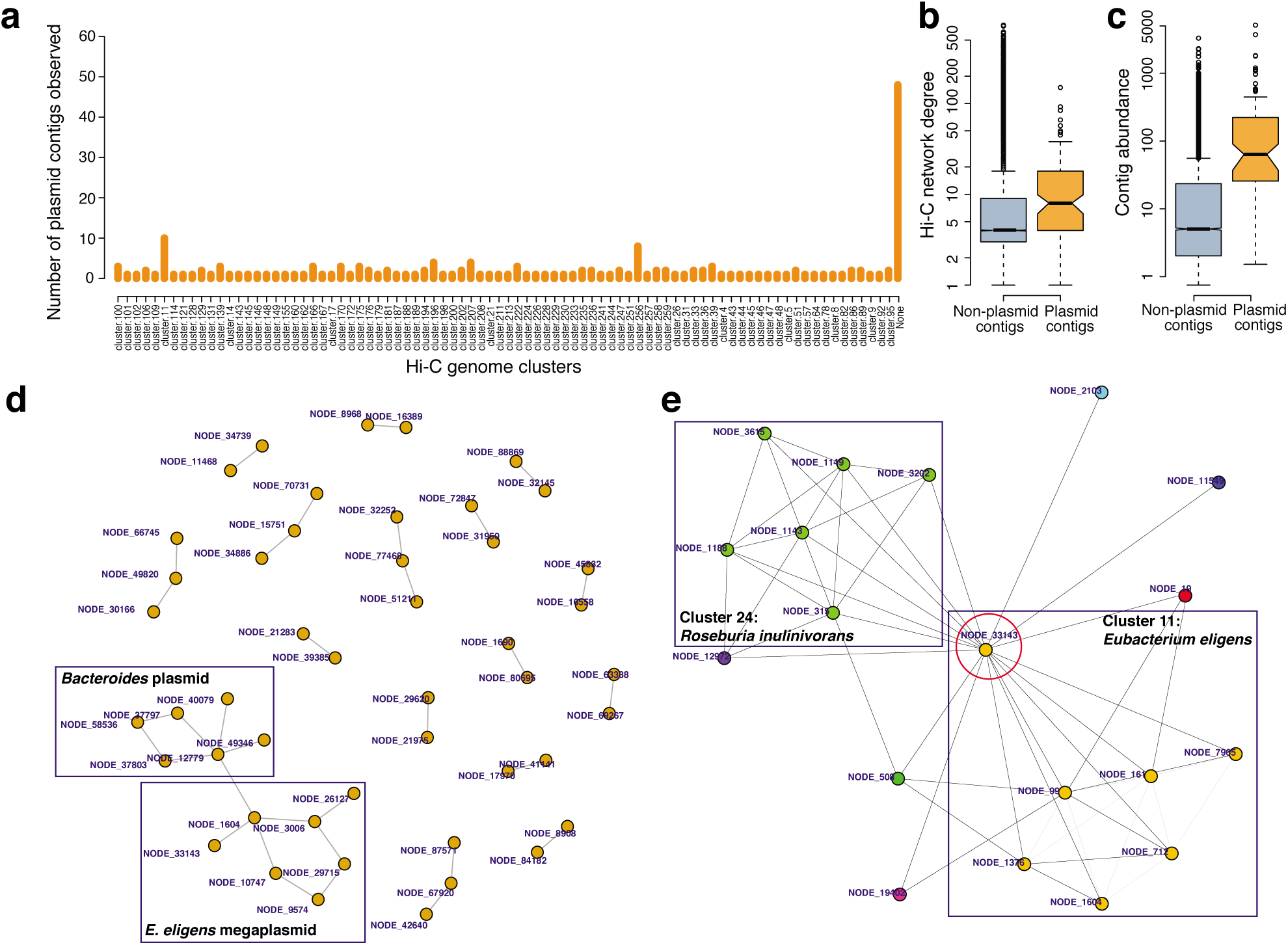
ProxiMeta associates plasmids with prokaryotic genomes and associates contigs from the same plasmid. (a) All contigs homologous to plasmid sequences with mapped Hi-C reads were separated by genome cluster, or assigned to a #x0022None“category (far right). (b) The degree of each clustered contig in the Hi-C graph (number of other contigs to which it is connected by Hi-C reads) according to classification status. (c) The abundance of each clustered contig in the Hi-C graph, as estimated by shotgun read pseudoalignment. (d) The Hi-C graph, restricted to plasmid contigs connected to at least one other plasmid contig. Pertinent features of the graph are indicated. (e) A subgraph of the Hi-C graph showing interactions of a plasmid contig (red circle) with two different highly complete high-quality genome clusters.

The genome cluster with the most plasmid contigs was cluster 11, which we inferred to be *Eubacterium eligens* based on k-mer hashes (Table S2). Notably, the reference genome for this strain carries a megaplasmid of approximately 600 kb in size. We confirmed that all of the plasmid contigs in this draft genome are homologous to this megaplasmid (Table S6), comprising a total of *∼*100 kb of contig sequence. However, other sequences similar to this megaplasmid were clustered into other genomes (in total comprising another*∼*100 kb, almost all in clostridial clusters; Table S7), suggesting that the megaplasmid might exist in multiple genomes, or that some related organisms carry different variants of this large plasmid. Moreover, given the high rate of gene gain and loss in bacteria (Kuo and Ochman, 2009; Mira, Ochman, and Moran, 2001; Puigbò et al., 2014), it is probable that the reference sequence is not definitive.

We next examined whether plasmid contigs exhibited specific patterns in the Hi-C connectivity graph. We observed that plasmid contigs generally had many more Hi-C read links than non-plasmid contigs (Figure 5b); potential explanations for this observation could include both high abundance of plasmid DNA relative to chromosomal DNA and gene sharing, such that plasmids are connected to multiple genomes. By examining mapped read-depths in the shotgun data, we found that plasmid contigs were, on average, much more abundant than those contigs without plasmid homology (Figure 5c).

Next, we grouped plasmid sequences together into putative plasmids, naively considering simple connected components in the Hi-C graph (Figure 5d). Different contigs in these plasmid clusters mapped to the same reference plasmid 62% of the time, strongly suggesting that this procedure identified and clustered plasmids (Table S7). The largest of these assemblies included sequences of the *E. eligens* megaplasmid mentioned above, though it was joined by a single apparently spurious link to contigs of a large *Bacteroidetes* plasmid.

To evaluate the evidence for gene sharing of plasmid contigs, we examined their connectivities to different genome clusters. Notably, a clustering assignment by ProxiMeta may reflect only a dominant host, rather than reflecting the full spectrum of Hi-C interactions of a plasmid contig. We therefore examined the local neighborhoods of the Hi-C graph around plasmid contigs nodes, and found several such instances where plasmids were linked to multiple clusters, suggesting gene sharing (Figure 5e, Figure S9). For example, contigs of the *E. eligens*-associated megaplasmid are connected to several complete genome clusters of *Clostridiales* species (Figure 5e, Figure S9), suggesting that in spite of their relatively divergent chromosomal genomes these species may yet share plasmid DNA with each other. Notably, average nucleotide identity (ANI) estimates for these genomes (*∼*70% for clusters 11 and 24) are low enough that highly conserved vertical inheritance is an unlikely explanation for these contacts, as in this analysis we require perfect alignments for Hi-C read mapping. This information about gene sharing can be exploited in future development of ProxiMeta to better understand this important component of microbial community dynamics.

## Discussion

The superior performance of ProxiMeta Hi-C relative to genome binning methods can be attributed to its direct ascertainment of physical proximity between DNA sequences. This physical, intra-cellular proximity represents a higher standard of evidence than indirect correlates such as sequence composition or estimated abundance, which are the two measures most commonly used by metagenome binning methods (Imelfort et al., 2014; Dick et al., 2009; Kang et al., 2015; Wu, Simmons, and Singer, 2016). In this paper we found the performance of ProxiMeta superior to a traditional binning approach as evaluated by standard metagenome quality metrics. Future improvements in the wet-lab and computational aspects of ProxiMeta will further improve the recovery of high-quality genomes from microbiome research efforts.

Another feature of ProxiMeta that is particularly valuable is its ability to localize specific sequences to particular genomes with high confidence. We demonstrate this feature in this study by linking plasmids to host genomes, but in principle the same could be applied to antibiotic resistance genes, secretion systems, phage, metabolic pathways, or any other sequences of interest. We anticipate that this technology will be of broad usefulness in medical, industrial, and academic applications.

One further important feature of ProxiMeta is that even small, incomplete genome fragments generated by this method represent relatively high-quality groupings, as they are based on physical proximity rather than indirect correlates. The large number of such high-quality fragments argues that further methodological improvements to ProxiMeta, or simply deeper Hi-C sequencing, have the potential to yield even higher numbers of high-quality genomes.

In conclusion, directly measuring physical co-occurrence of DNA sequences within cells makes ProxiMeta an effective method for the characterization of complex microbial communities, both in the deconvolution of complete genomes and the pursuit of targeted biological questions.

## Methods

### Sample preparation

A fecal sample was collected from a human subject. For shotgun sequencing, DNA was extracted using the zymoBIOMICS DNA Mini kit and 100 ng was sheared to 500 bp average insert length and used to create a shotgun library using the HyperPrep kit (KAPA Biosystems). Approximately 200 *μ*L of solid material from the same sample was crosslinked for Hi-C using standard protocols (Burton et al., 2014) and split into two fractions. Each fraction was used to generate a Hi-C library using the proprietary ProxiMeta protocol developed by Phase Genomics (standard Hi-C protocols can be found in (Burton et al., 2014; Belaghzal, Dekker, and Gibcus, 2017)). One Hi-C sample was fragmented using Sau3AI (New England Biolabs) and the other using MluCI (New England Biolabs) prior to proximity-ligation.

The shotgun and Hi-C libraries were sequenced on the Illumina HiSeqX platform, generating 151 bp paired-end reads. Sequencing of the shotgun library produced 250,884,672 read pairs. Sequencing of the Hi-C libraries generated 41,733,770 read pairs for the Sau3AI library and 48,798,091 read pairs for the MluCI library.

### Hi-C read processing

Using the Hi-C read datasets generated as described above, we trimmed each read to 75 bp to avoid discarding reads sequencing through a Hi-C junction. We mapped each read dataset (MluCI and Sau3AI) to the shotgun assembly described above using bwa aln (Li and Durbin, 2009) while requiring perfect matches (option -n 0). A total of 907,243 read pairs from the MluCI dataset and 1,361,063 read pairs from the Sau3AI dataset mapped mates to different contigs, and these reads were used for deconvolution.

### Metagenomic assembly

We trimmed adapter sequences from shotgun reads using BBDuk (*BBTools*) with options k=23, ktrim=r, mink=12, hdist=1, minlength=50, –tpe, –tbo. Next, we performed quality trimming of the reads using BB-Duk and options qtrim=rl, trimq=10, minlength=50, chastityfilter=True. We then normalized read coverage using BBNorm with options target=40, mindepth=2. With this trimmed and normalized dataset, we performed a *de novo* shotgun assembly using metaSPAdes and default parameters (Nurk et al., 2017). We finally evaluated this assembly (and downstream groupings) using quast (Mikheenko, Saveliev, and Gurevich, 2016).

### ProxiMeta deconvolution

We performed deconvolution of the shotgun assembly into genomes using ProxiMeta software, which is similar to the previously described MetaPhase technique (Burton et al., 2014), but using proprietary clustering and post-processing steps tooled towards high-complexity samples. We filtered out all reads that were not properly paired, unmapped, non-uniquely mapped, had a MAPQ score less than 20, or were paired with a mate with an identical seqid. We filtered out contigs that were less than 1000 bp in size, or which contained fewer than 2 restriction sites for the relevant enzyme. We combined datasets of the two restriction enzymes into a graph, and applied a normalization to the read counts connecting each pair of clusters by accounting for the estimated abundance of each contig. We clustered contigs into genome clusters using a proprietary MCMC-based algorithm based on their Hi-C linkages. This analysis yielded a total of 252 genome clusters.

### Genome binning

To generate an independent set of genome bins to which we could compare ProxiMeta results, we used the same shotgun assembly and MaxBin v2.2.4 (Wu et al., 2014; Wu, Simmons, and Singer, 2016) to group contigs into genome bins, while discarding contigs less than 1Kbp (MaxBin default), maximum Expectation-Maximization iteration number of 50, and probability threshold of EM final classification of 0.9. This yielded a total of 189 genome bins.

### Analysis of genome clusters and bins

We evaluated each genome cluster using a variety of tools. We used checkM (Parks et al., 2015) to estimate the likely completeness and contamination of each genome cluster based on marker genes, including a taxonomic assignment according to pplacer (Matsen, Kodner, and Armbrust, 2010). We used Mash (Ondov et al., 2016) to search a RefSeq database for similar reference genomes using k-mer hashing. We further used Infernal (Nawrocki and Eddy, 2013) with the Rfam database (Nawrocki et al., 2015) (with options–rfam –noali –cpu=2) and Aragorn (Laslett and Canback, 2004) (with options -ps -w) to detect rRNA and tRNA genes respectively in genome clusters.

For comparisons of ProxiMeta-derived and binning-derived genomes to closely-related reference genomes, we used MetaQuast with default parameters (Mikheenko, Saveliev, and Gurevich, 2016) to generate similarity summaries and Circoletto (Darzentas, 2010) to visualize similarity comparisons based on BLAST identity. For genomes generated by MaxBin, we selected genome bins in a similar fashion to those selected from ProxiMeta clusters, though we were obliged to loosen the inclusion criteria slightly to >97% complete, <2% contamination, and >580/1000 k-mer hashes to the reference.

### Abundance quantification

We quantified abundance of each contig in each of the two genome groupings using pseudoalignment of the shotgun reads with kallisto using default parameters (Schaeffer et al., 2017; Bray et al., 2016). For each cluster, we used the median abundance (quantified as a transcripts per million estimate) among all component clusters as a point estimate of the overall cluster abundance. We preferred the median because some contigs (particularly small ones) showed dramatically higher estimates, possibly attributable to higher copy-number elements such as plasmids or to collapsed multi-copy regions, yielding a long upper tail of the abundance distribution.

### Plasmid analysis

We downloaded all sequences labeled as plasmids from NCBI (accessed August 18, 2017) and subjected these to light manual curation to remove duplicates and sequences smaller than 500 bp. We used BLAT (Kent, 2002) to search all contigs of the raw shot-gun assembly against this database (options -t=dna-q=dna -maxIntron=0 -minIdentity=95 -minMatch=5-minScore=200). We further filtered out all alignments shorter than 500bp or covering <10% of the contig’s length to remove spurious alignments and contigs of which plasmid sequences were only a very small proportion. We took the remaining alignments as evidence that contigs contained plasmid sequence. For further analysis we used the igraph R library (Csardi and Nepusz, 2006) to conduct and visualize network analyses of the Hi-C connectivity graph.

## Data availability

Short-read data has been deposited into NCBI under the BioProject ID: PRJNA413092 with short reads found under SRR6131124, SRR6131123 and SRR6131122.

